# Ncl1 mediated metabolic rewiring critical during metabolic stress

**DOI:** 10.1101/529578

**Authors:** Ajay Bhat, Rahul Chakraborty, Khushboo Adlakha, Ganesh Agam, Kausik Chakraborty, Shantanu Sengupta

## Abstract

Nutritional limitation has been vastly studied, however, there is limited knowledge of how cells maintain homeostasis in excess nutrients. In this study, using yeast as a model system, we show that some amino acids are toxic at higher concentrations. With cysteine as a physiologically relevant example, we delineated the pathways/processes that are altered and those that are involved in survival in presence of elevated levels of this amino acid. Using proteomics and metabolomics approach, we found that cysteine upregulates proteins involved in amino acid metabolism, alters amino acid levels, and inhibits protein translation, events that are rescued by leucine supplementation. Through a comprehensive genetic screen we show that leucine mediated effect depends on a tRNA methyltransferase (Ncl1), absence of which decouples cell’s transcription and translation, inhibits the conversation of leucine to ketoisocaproate and leads to TCA cycle block. We therefore, propose a role of Ncl1 in regulating metabolic homeostasis through translational control.

## Introduction

Cell requires optimal nutrients for the synthesis of macromolecules like lipid, protein and nucleotides. Activation of these anabolic processes and the repression of catabolic process like autophagy promote cellular growth. On the other hand, when nutrients are limiting, anabolic processes are inhibited, and autophagy is activated ^1^. Notably, among these processes, protein synthesis consumes a large portion of nutrients and energy ^2 3^. Amino acids are the building block of proteins, and therefore its availability regulates the process of translation ^4^. Insufficient amino acid levels leads to the inhibition of protein synthesis ^5^ and induce the expression of genes required for synthesis of amino acids ^6^. Eukaryotic cell have two well-known pathways for sensing amino acids, which includes target of rapamycin complex 1 (TORC1) ^7^ and general control nonderepressible 2 (Gcn2) ^8^. Limitation of amino acids promotes Gcn2 mediated signalling ^9^, and inhibits TORC1 ^10^, both of which lead to inhibition of protein synthesis ^1 11^.

In contrast, there are reports indicating that excess of amino acids can lead to cellular toxicity ^12 13^. Elevated branch chain amino acids like Leu, Ile and Val are associated with insulin resistance ^14 15^ and is also responsible for neurological damage ^16^. Metabolic profiling has demonstrated that high levels of aromatic amino acids could be associated with the risk of developing diabetes ^15 17^. Increased levels of Phenylalanine either due to genetic mutation or exogenous supplementation leads to abnormal brain development ^18 19^. Glutamate mediated excitotoxicity leads to neuronal death ^20^, neonatal mice injected with high doses of glutamate shows increases in body fat and damaged hypothalamus region ^21^. Increased Histidine intake in animals resulted in hyperlipidemia, hypercholesterolemia, and enlarged liver ^12 22^. High levels of sulfur amino acids, Cysteine and Methionine are reported to be toxic in various model systems ^23 24 25 26 27^. Supplementation of excessive Methionine in rats supresses food intake, diminishes growth and also induces liver damage ^28 12^. In mammals, excessive levels of Cys have been demonstrated to be neurotoxic in many in-vivo and in-vitro studies ^29 30^. Reports also suggest that elevated levels of cysteine may be associated with cardiovascular disease ^31^, ^32^.

Cellular response due to nutrient deprived conditions, and how nutritional deficiency is sensed by the cell has been well studied. However, information of how cell responds and survives during nutritional excess is lacking. This information will help us understand how cells maintain homeostasis when it accumulates excess of an amino acid, which is important since consumption of single amino acids as health supplements, and their use as flavoring agents has increased in recent years.

In this study, using *Saccharomyces cerevisiae as a model*, we used a combination of genetic, proteomic, transcriptomic and metabolic approach to understand cysteine induced systemic alteration. We show that cysteine inhibits protein translation and alters the amino acid metabolism, effects which are reversed by supplementation of leucine. We also show that a tRNA methyltransferase (NCL1) is involved in survival during cysteine stress, absence of which inhibited the conversation of leucine to ketoisocaproate (KIC), a necessary step for mitigating the effect of cysteine. Thus, this study not only uncovered the cellular insights during high levels cysteine, but also highlighted the novel role of NCL1 in regulating the metabolism during excess cysteine.

## Results

### Metabolic alterations due to excess amino acids induce growth defect

Growth screening of *S cerevisiae* (BY4741) in the presence of high concentrations of amino acids revealed that a few amino acids - cysteine, isoleucine, valine, tryptophan and phenylalanine inhibited growth (Fig. 1A). To test if the imbalance due to excess of an amino acid resulting in the growth inhibition could be abrogated in the presence of other amino acids, we did a comprehensive amino acid supplementation screening. We found that the amino acid, leucine could completely rescue the growth inhibition due to isoleucine, valine, tryptophan and phenylalanine but not cysteine, where the rescue was partial (Fig 1B). The effect of leucine could be because the yeast strain used in this study, BY4741, is auxotrophic for leucine. To confirm this, we performed the same experiment using a prototrophic wild type strain, S288C, and found that except cysteine, none of the other amino acids inhibited growth (Fig 1C). Further, the growth inhibitory effect of cysteine was lower compared to BY4741. This suggests that intracellular concentration of leucine might play a role in alleviating amino-acid mediated toxicity. This was further confirmed by deleting the gene involved in leucine biosynthesis (LEU2) in a prototrophic strain (S288C), which increased the sensitivity of this strain towards cysteine (Fig 1D) and other amino acids (Fig 1E). Since, cysteine was the only amino acid whose growth inhibitory effect could not be completely recovered by leucine, we focussed our attention in understanding the effect of cysteine in yeast.

**Fig 1.**
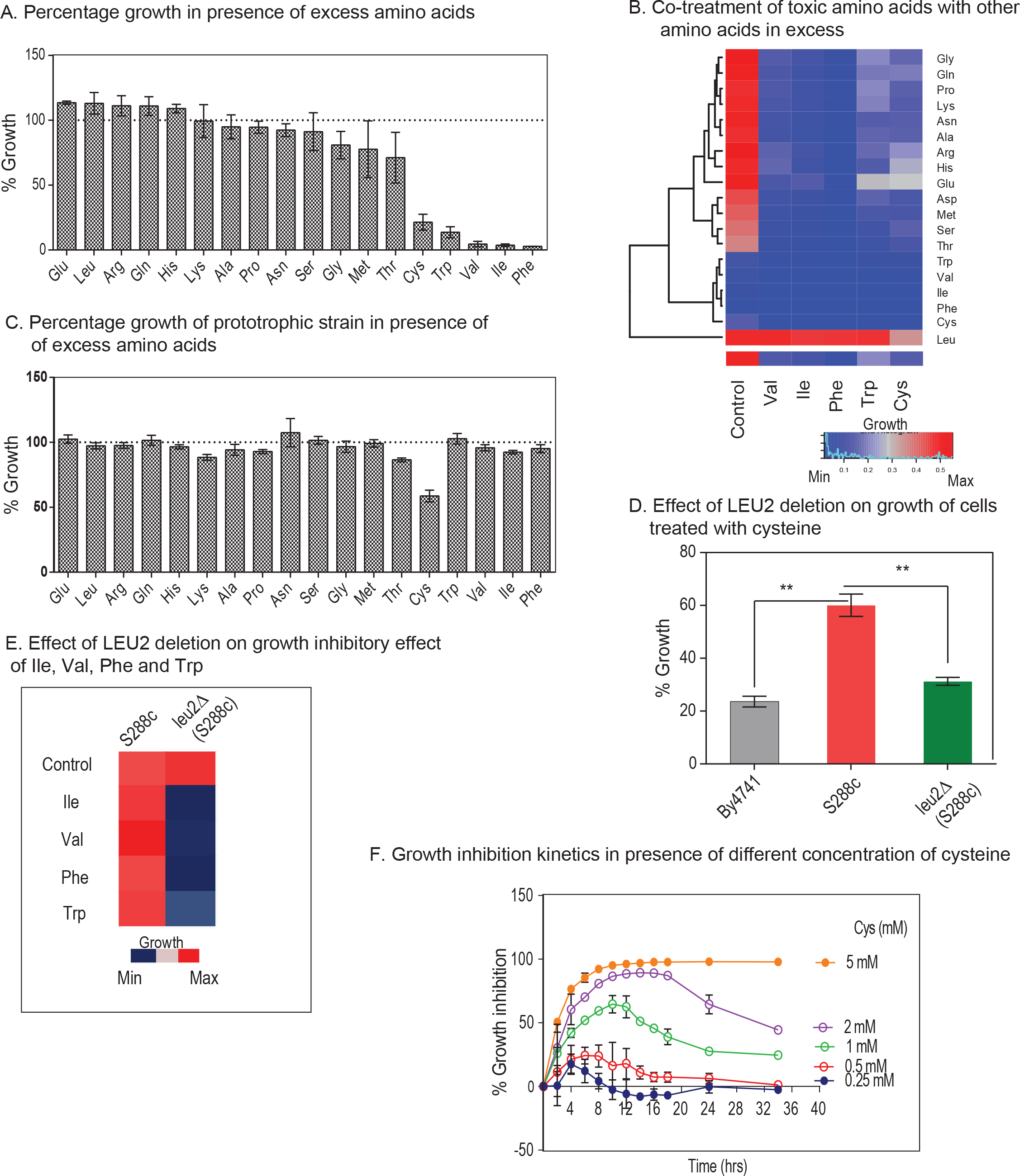
Metabolic alterations due to excessive amino acids induces growth defect. A. Cells of BY4741 strain were grown in higher concentration of amino acids (~10 times higher the concentrations of amino acids than the recommended in the media) for 12 hrs and growth was monitored by measuring O.D at 600 nm. Percentage growth was then calculated with respect to the untreated cells (growth in SC media containing basal concentration of amino acids). B. The higher concentration of toxic amino acids (Cys, Trp, Val, Ile, Phe) were co-supplemented with higher levels (10 times as that of in media) of other amino acids and growth was measured at 600 nm using multimode reader. The figure represents the growth matrix of cells grown with different combination of amino acids. The bottom most row represents the growth of the cells during normal SC media (Control) and treatment with high doses of single amino acids (Val, Ile, Phe, Trp and Cys). Growth of the cells during these conditions co-treated with other amino acids (at higher concentrations) is represented in the above matrix. C. Prototrophic strain (S288C) was grown with higher concentration of different amino acids for 12 hrs and then O.D (at 600 nm) was measured to calculate the percentage growth with respect to untreated cells (growth in basal SC media). D. Percentage of growth during cysteine treatment (1 mM) in BY4741, S288C and leu2Δ (LEU2 deleted in S288C background) strains. E. Growth of S288C and leu2Δ (LEU2 deleted in S288c background) strains in presence of toxic amino acids (Ile, Val, Phe, and Trp) F. Overnight grown cultures of yeast was re-inoculated at an O.D of 0.1 in SC media and growth was monitored in presence (0.25 mM - 5 mM) and absence of L-cysteine at different time point. The graph shows the kinetics of growth inhibition due to different concentrations of cysteine. Percentage growth inhibition was calculated by using formula {(O.D of control cells – O.D of cysteine treated cells)/O.D of control cell x 100}.

Cysteine inhibits the growth of yeast (BY4741) in a dose dependent manner (Fig S1A). Interestingly, for each concentration of cysteine (other than 5 mM, at which there was very minimal growth), the growth inhibition increased with time, reached maxima, and then decreased (Fig. 1F). The recovery in the growth inhibition clearly indicates that cells have response mechanisms that lead to cellular adaptation.

### Amino acid metabolism and protein translation plays a vital role in cysteine toxicity

To understand the pathways altered due to cysteine treatment we used a iTRAQ-based proteomics approach and identified proteins that were differentially expressed at 6 hours (early time point) and 12 hours (later time point) (Fig S2A, and Supp Table 1 & 2). We found that proteins involved in amino acid metabolism, were mostly upregulated at both the time points (Fig 2A). Notably, in line with the genetic links to leucine metabolism, there was a strong induction in the enzymes linked to the synthesis of branched chain amino acids including Leu, Ile, Val. However, exogenous addition of leucine reverted the expression of these metabolic enzymes, suggesting that induction of these enzymes was the response towards cysteine induced stress (Fig 2B).

**Fig 2.**
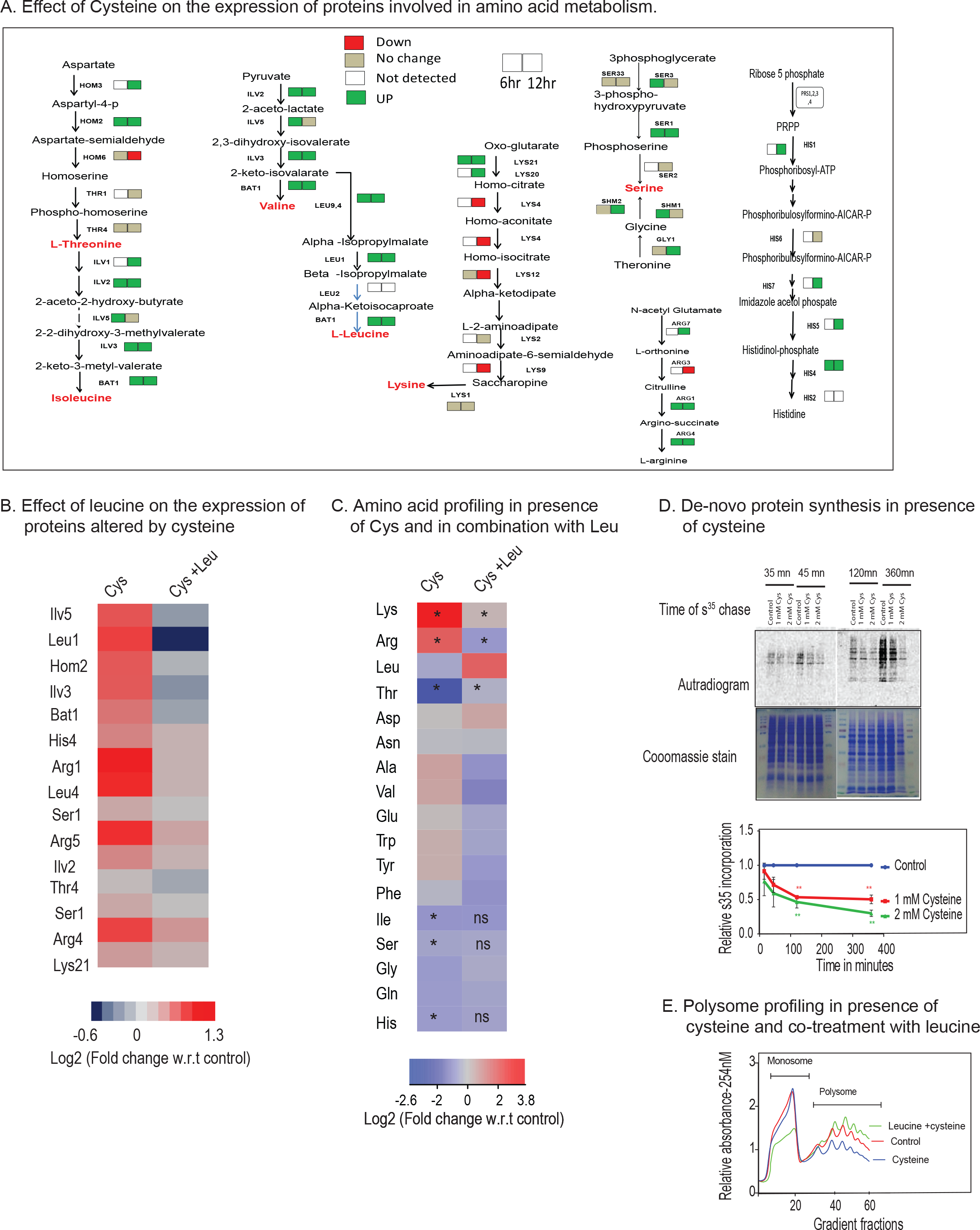
Amino acid metabolism and protein translation plays a vital role in cysteine toxicity. A. Cells were treated with cysteine for 6 and 12 hrs, and then iTRAQ based relative quantification of proteins was performed. The figure represents the directionality of the expression of proteins involved in amino acid metabolism during 6 and 12 hrs of cysteine treatment. Left side and right side box for each protein denotes its expression at 6 and 12 hrs of cysteine treatment, respectively. Red, green, beige colored box represents the proteins downregulated, upregulated and not changed, respectively due to cysteine treatment. However, white colored box represents the proteins not detected in the proteomics experiment. B. Heat map represents the relative expression of proteins (with respect to untreated condition) involved in amino acid metabolism during cysteine treatment and their status during co-treatment of leucine and cysteine. C. Heat map represents the log fold change of different amino acids during cysteine (Cys) treatment and combination of leucine and cysteine (Leu+Cys). In the first column the amino acids significantly altered by cysteine are labelled by “*” (p <0.05), however if the levels of these amino acids are significantly reverted by co-treatment of leucine and cysteine then in the second column they are labelled by “*” (p <0.05) or otherwise “ns” (non-significant). D. S^35^-Methionine incorporation kinetics in presence of different concentration (1 mM and 2 mM) of cysteine for different time points. In the upper panel, the upper blot is for autoradiogram, which signifies S^35^-Met incorporation in proteins and the lower blot represents coomassie stain of the same proteins, which represents equal loading of proteins. The lower panel is the quantification of S^35^ incorporation with respect to untreated cells. *“**” indicates p < 0.01.* E. Polysome profiling representing monosome and polysome fractions in presence and absence of cysteine treatment and during co-treatment of leucine and cysteine.

We therefore, determined the relative levels of amino acids in wild type and cysteine treated cells and found that the levels of lysine was extremely high while that of threonine extremely low in the presence of cysteine (Fig 2C). The levels of other amino acids were also altered *albeit* to a much lesser extent. Furthermore, cysteine-induced increase in Lys, Arg and decrease in Thr was completely reversed in the presence of leucine.

Proteomics data also revealed alteration in proteins involved in translation machinery (Fig S2A) and since leucine is known to activate protein translation, we tested if cysteine toxicity is associated with translation arrest. For this, we performed S^35^-Methionine incorporation assay to analyse the effect of cysteine on *de-novo* protein synthesis and found that cysteine leads to a significant reduction in the rate of incorporation of S^35^-Methionine compared to untreated cells (Fig 2D). To prove that this effect was not due to intracellular conversion of cysteine to methionine through homocysteine, we did the same experiment in met6Δ strain (which blocks the conversion of homocysteine to methionine). Even in this strain, cysteine inhibited the incorporation of S^35^-Methionine (Fig S2B). Translation inhibition was also confirmed using polysome profiling, where we found that high levels of cysteine decrease the abundance of polysomes (Fig 2E). Co-treatment of leucine and cysteine drastically increase the levels of polysomes, and thus maintains the active translational state of the cells. This clearly shows that cysteine induced amino acid alterations and translational defect could be rescued by leucine treatment.

### Genetic interactors of leucine mediated rescue

To better understand the mechanism of leucine induced alleviation of cysteine toxicity, we performed a genome wide screen to identify the genetic interactors involved in leucine mediated abrogation of cysteine induced toxicity. An initial screen for cysteine sensitivity revealed 50 deletion strains to be sensitive (Fig 3A and Supp. Table 3). Deletion of genes involved in amino acid metabolism like THR4 (threonine synthase), THR1 (Homoserine kinase), HOM3 (Aspartate kinase), LEU3 (Regulates branched chain amino acid synthesis), STP2 (Regulates the expression of amino acid permeases) and YML082W (paralog of STR2) were among these sensitive strains, which further highlighted the significance of amino acid metabolism during high levels of cysteine.

**Fig 3.**
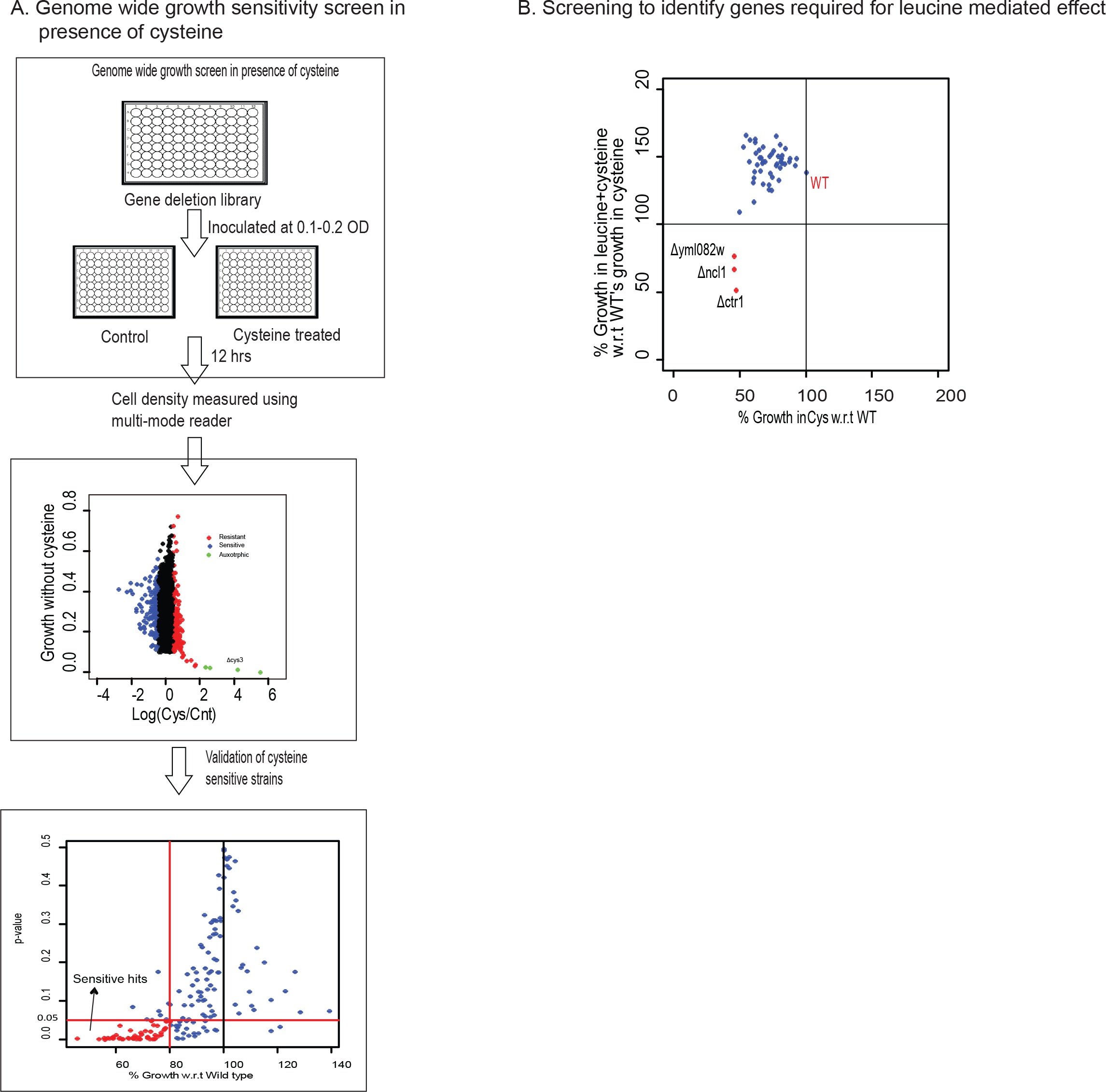
Genetic interactors of leucine mediated rescue. A. Whole genome screen in presence of 1mM cysteine. Upper most panel represents the schematic of the genome wide screen. The middle panel represents scatter plot between logarithmic ratio of growth in presence and absence of cysteine versus growth in basal media (SC media), for each deletion strain. In this panel the deletion strains, which show growth sensitivity and resistance for cysteine, are symbolized with blue and red dots, respectively. Strains which were very slow growing and show growth advantage with cysteine as denoted by green dots. Lower most panel represents the validation of the cysteine sensitive strains. Strains which show growth sensitivity towards high levels of cysteine in first round of whole genome screen were again grown in presence of cysteine and growth was monitored and compared with Wild type (Wt) strain. Three biological replicates were performed and average percentage growth of each strain with respect to Wt was calculated. Deletion strains whose growth was significantly different from Wt (p-value ≤ 0.05), and grows atleast 20% less than Wt, were considered to be sensitive for cysteine, and are denoted by red dots. B. Screening to identify genes required by leucine for alleviating cysteine induced toxicity. Cysteine sensitive deletions strains were grown with and without higher concentration of cysteine (1 mM), leucine (5 mM) and combination of leucine and cysteine. Figure represents the scatter plot between percentage growth during cysteine treatment versus percentage growth during combination of leucine and cysteine treatment, with respect to wild type’s growth during cysteine treatment. Deletion strains in the below left quadrant, which are denoted by red dots, represents the strains which are sensitive to cysteine but are not recovered by leucine.

We further checked the growth of all these sensitive strains in the presence of cysteine and leucine. A reversal of cysteine induced toxicity was observed in almost all these deletion strains, except for three strains, yml082wΔ, ncl1Δ and ctr1Δ (Fig 3B). YML082W is an uncharacterized gene, which is predicted to have carbon sulphur lyase activity, and is a paralog of cystathionine gamma synthase (str2), which converts cysteine to cystathionine. CTR1 is a high affinity copper transporter of plasma membrane ^33^ and has a human homolog SLC31A1 ^34^. NCL1, is a SAM dependent m5C-methyltransferase, and methylates cytosine to m5C at several positions in various tRNA ^35 36^. Interestingly, a leucine tRNA (CAA), is the only tRNA where it methylates at the wobble position ^35 37^. This coupled with the fact that cysteine mediated amino acid alterations are reverted by leucine treatment led us to focus on the role of NCL1 in abrogating the growth inhibitory effect of cysteine.

### Leucine reprograms proteome through NCl1 to recover cysteine-mediated toxicity

We performed growth kinetics to confirm the role of NCL1 during cysteine induced growth inhibition and leucine mediated recovery. Deletion of NCl1 makes the cells incapable of abrogating cysteine induced toxicity even in the presence of leucine (Fig 4A). To confirm the Ncl1 dependent effect of leucine, we analysed the proteomic profile of Wt and ncl1Δ strain in the presence of cysteine alone and a combination of cysteine and leucine. While cysteine treatment in Wt cells lead to the upregulation of proteins involved in amino acid metabolism both at 6 and 12 hours and translation machinery at 12 hours, in ncl1Δ strain it leads to down-regulation of proteins involved in translational machinery and amino acid metabolism (Fig S3A, Supp. Table 4). This indicates that NCL1 may play a central role in regulating translation and metabolic rewiring that is required to alleviate cysteine toxicity. In Wt cells, the cysteine induced differentially expressed proteins were significantly reverted by leucine treatment (Fig 4B, Fig S3B). Proteins upregulated by cysteine in Wt were less upregulated in ncl1Δ during cysteine treatment, and were unaltered during leucine treatment. This indicates global down-regulation of cysteine induced response at the protein level in ncl1Δ, which could even not be reverted by leucine treatment. To confirm if the changes induced by NCl1 in the presence of cysteine is at the transcriptional or translational level, we quantified changes in mRNA and protein levels during cysteine stress in Wt and ncl1Δ cells (Supp. Table 5). In Wt, we observed correlated changes in mRNA and protein expression upon cysteine treatment (rho=0.3, p<0.0001, Fig. 4C top panel), however, no such correlation was found in Δncl1 strain (rho= 0.07, p=0.18, Fig.4C bottom panel). This indicates a decoupling of translation and transcription in Δncl1 strain during cysteine treatment.

**Fig 4.**
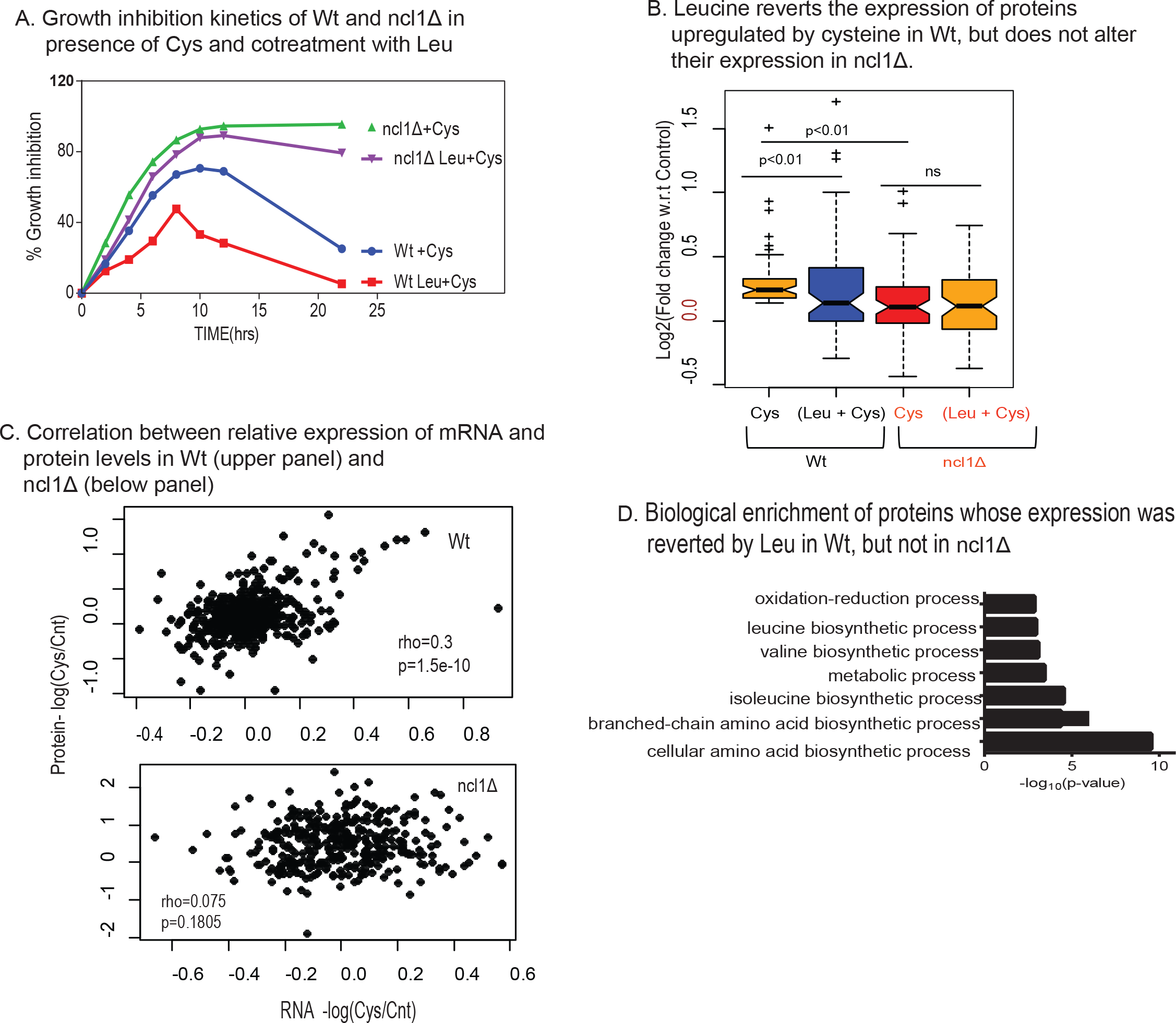
Leucine reprograms proteome through NCl1 to recover cysteine-mediated toxicity. A. Percentage growth inhibition of wild type (Wt) and ncl1Δ strain, in presence of cysteine and combination of leucine and cysteine. B. Box plot represents the relative expression of proteins during co-treatment of leucine and cysteine in Wt and ncl1Δ, for proteins which were upregulated by cysteine in Wt. C. Spearman rank correlation between relative expression of mRNA (obtained from RNA-seq) and protein levels (from iTRAQ based relative proteomics) in Wt (upper panel) and ncl1Δ (lower panel) D. The bar graph represents the biological pathways enriched from proteins whose expression was reverted during co-treatment of leucine and cysteine in Wt, but not to the similar extent in Δncl1. Classification was done using Database for Annotation, Visualization & Integrated Discovery (DAVID), and pathways with p-value ≤ 0.05 are plotted in the figure.

To identify the pathways that could potentially be affected by NCL1 deletion, we analyzed the proteins whose expression was altered in cysteine and reverted by leucine in Wt but not in NCL1 deleted cells. These proteins belong to amino acid metabolism, specially branched chain amino acid biosynthetic proteins and oxidation-reduction process (Fig 4D). This indicates that NCL1 might be necessary for leucine to abrogate the amino acid imbalance induced by cysteine.

### Metabolism of leucine via NCL1 plays a vital role during cysteine induced stress

To understand the role of NCL1 at the metabolic level, we measured the amino acids in ncl1Δ cells during cysteine treatment and co-treatment of cysteine with leucine. Interestingly, in ncl1Δ cells, the amino acid profile of cysteine treated cells was similar to cells co-treated with cysteine and leucine as evident from their strong correlation (r^2^ = 0.99. p-value= 6.2 e-10, Fig 5A) This further suggests that leucine mediated metabolic rewiring depends on NCL1. Interestingly, we found a significant accumulation of leucine in ncl1Δ cells compared to Wt when these cells were treated either with leucine alone or in the presence of cysteine suggesting a defect in leucine metabolism in ncl1Δ strain (Fig 5B). Leucine is metabolized to alpha-ketoisocaproate (KIC) through branched chain aminotransferase (BAT) ^38^. The expression of BAT1 was about 2 fold lower in ncl1Δ strain than Wt in the presence of cysteine (Fig 5C) and the levels of KIC was markedly reduced in cysteine treated ncl1Δ strain even in the presence of leucine (Fig 5D). This confirms that the conversion of leucine to KIC is inhibited in ncl1Δ strain in the presence of cysteine which may explain the inability of leucine to abrogate cysteine induced growth defect in this strain. To confirm the role of KIC during cysteine treatment, we analysed the effect of KIC on growth in Wt and ncl1Δ strain. Interestingly, although leucine could not rescue cysteine toxicity in ncl1Δ strain, KIC could rescue the toxicity in both ncl1Δ and Wt strain (Fig 5E). This further proves that conversion of leucine to KIC is a necessary step for survival in the presence of cysteine. Interestingly, KIC was able to rescue the growth inhibition of other toxic amino acids similar to leucine (Fig. 5F).

**Fig 5.**
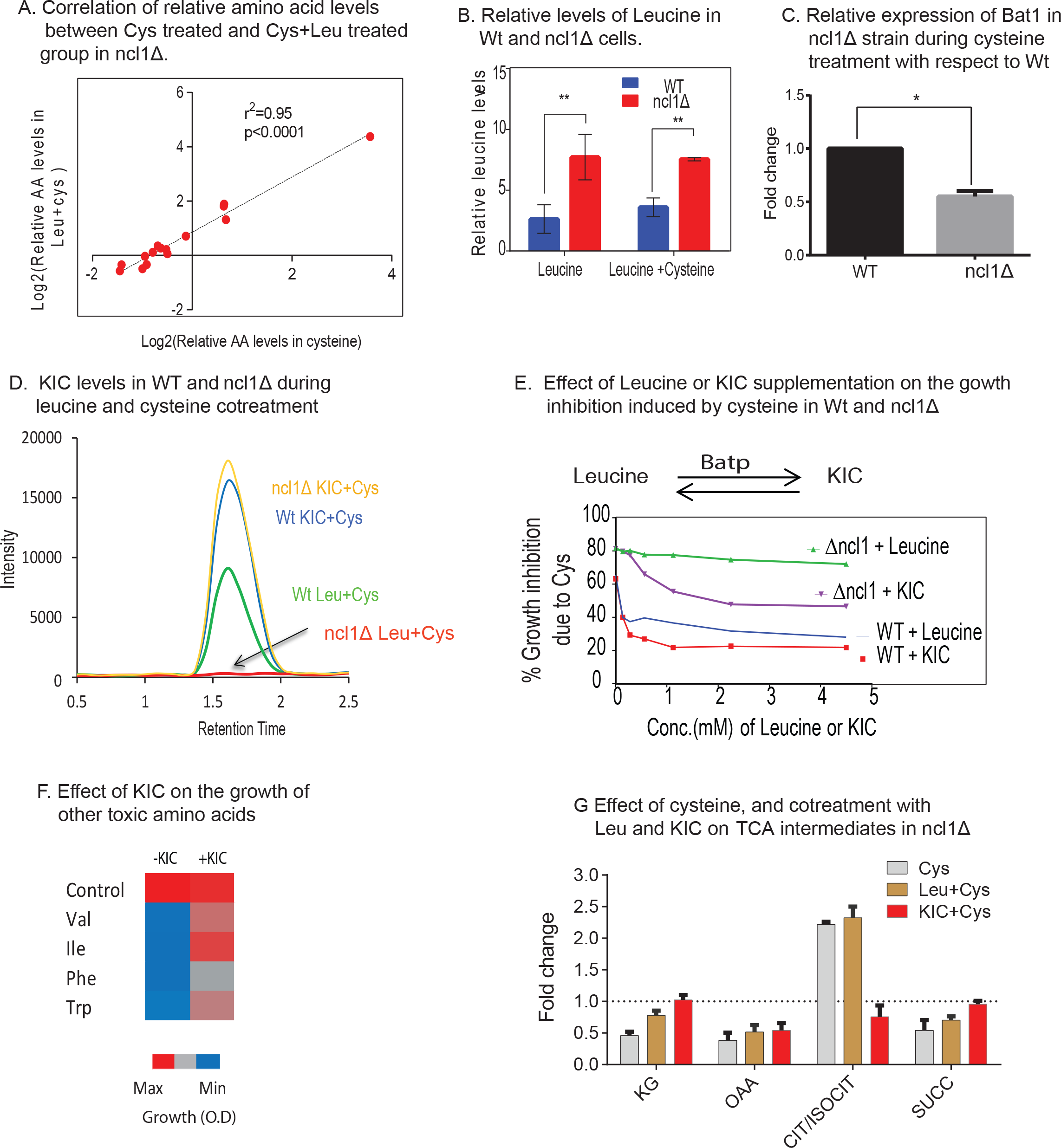
Metabolism of leucine via NCL1 plays a vital role during cysteine induced stress. A. Relative amino acid (AA) levels in ncl1Δ strain during Cys treatment and combination of Leu and Cys treatment were measured, and were analysed by calculating the correlation between these two groups. B. Bar graph represents the relative levels of leucine in Wt and ncl1Δ cells, during leucine and combination of leucine and cysteine, normalized with respect to levels of leucine in untreated Wt strain. *“**” indicates p<0.01.* C. Relative expression of BAT1 in Wt and ncl1Δ during cysteine treatment, normalized with respect to its expression in cysteine treated Wt. *“*” indicates p<0.05.* D. KIC levels were measured using MRM based LC-MS technique. The figure represents the overlapped peak intensity of a transition of KIC (121/101 m/z), measured from Wt and ncl1Δ cells during co-treatment of leucine and cysteine. KIC was also measured in both these cells during co-treated of KIC and cysteine. E. Cells were co-treated with cysteine (1 mM) and different concentration of Leucine or KIC, and then growth was measured after 12 hrs. The graph represents the growth inhibition of Wt and ncl1Δ strain due to cysteine in the basal media, and during its co-treatment with varying concentration of KIC and Leucine. F. Growth of cells in presence of toxic amino acid (Ile, Val, Phe and Trp), and during their respective co-treatment with KIC. G. TCA intermediates were measured by MRM based LC-MS technique. The bar plot represents their relative levels during cysteine treatment and its combination with KIC or Leucine in ncl1Δ strain.

Both KIC and Leucine have been shown to activate TOR ^39^, and Bat1 deletion leads to compromised TORC1 activity ^39^. Bat1 exhibits leucine dependent interactions with the TCA enzyme aconitase, and deletion of Bat1 leads to TCA cycle block ^39^. Thus, we measured the levels of TCA intermediates in ncl1Δ strain during cysteine treatment and its co-treatment with leucine and KIC. We found that cysteine treatment resulted in accumulation of citrate, with decreased levels of other TCA intermediates (alpha-ketoglutarate, succinate and oxaloacetate) (Fig. 5G). However, KIC but not leucine could reverse the levels of most of these metabolites. Citrate accumulation could potentially decrease the levels of pyruvate ^40^, which prompted us to check for the levels of pyruvate during cysteine treatment. We found that cysteine treatment significantly lowered the levels of pyruvate in ncl1Δ strain (Fig. 6A).

**Fig 6.**
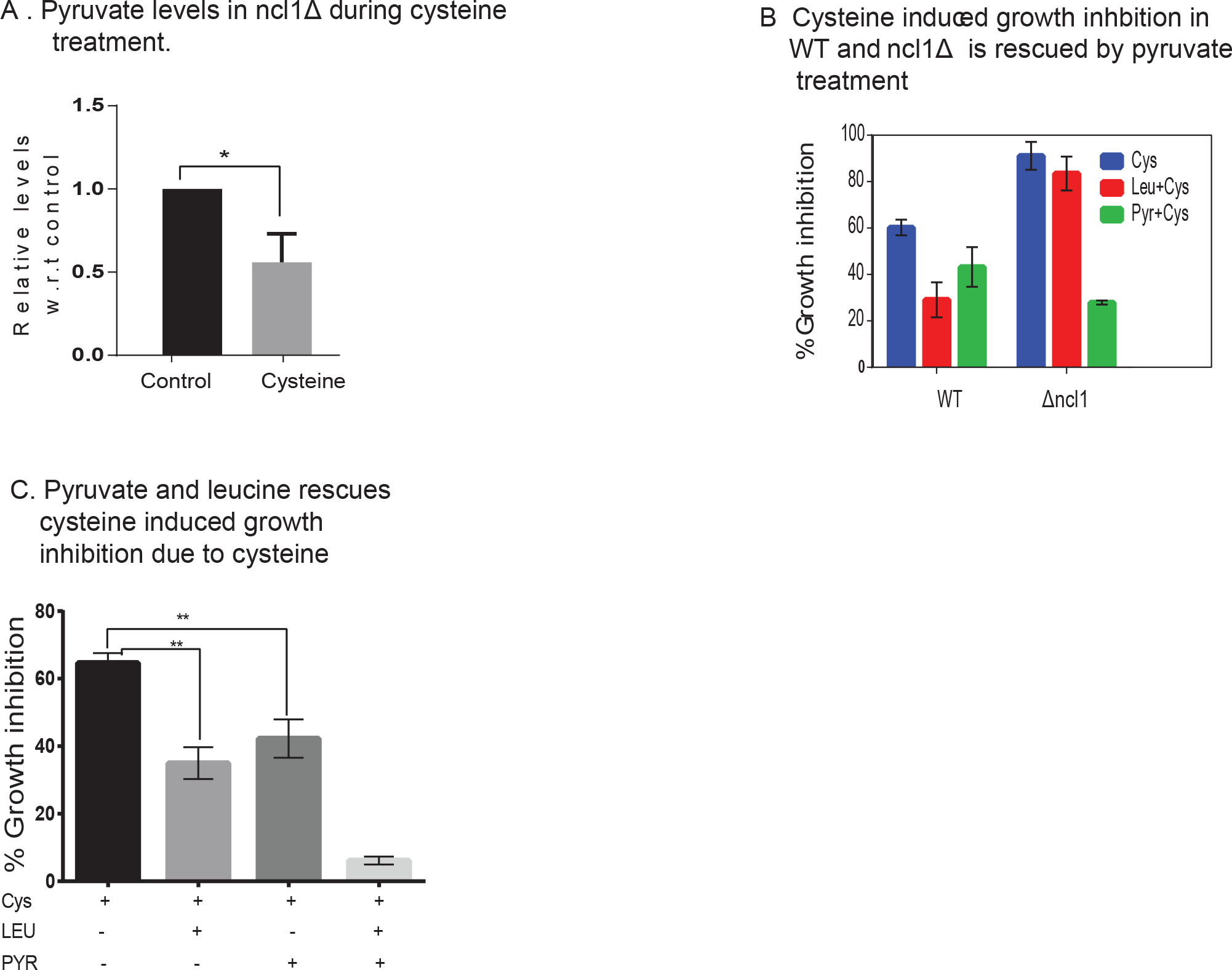
Pyruvate and leucine co-treatment completely rescues the toxicity of cysteine. A. Relative levels of pyruvate in ncl1Δ cells treated with cysteine, with respect to corresponding untreated control. “*” indicates p<0.05. B. Effect of pyruvate and leucine on the growth inhibition induced by cysteine in Wt and ncl1Δ cells. C. Growth inhibition of cysteine during cotreatment of leucine and pyruvate simultaneously. *“**” indicates p<0.01.*

To understand the significance of pyruvate levels, we measured the growth of Wt and ncl1Δ cells during exogenous addition of pyruvate. In Wt cells, pyruvate (5mM) was able to decrease the growth inhibition due to cysteine by about 35% (Fig 6B). Leucine at the same concentration was able to decrease the growth inhibition by about 45%. In ncl1Δ strain, where leucine could not abrogate the growth inhibition of cysteine, pyruvate could decrease the inhibition by more than 60%. Interestingly, we found that leucine and pyruvate had a synergistic effect and in combination could almost completely abrogate the cysteine induced inhibition (Fig. 6C), suggesting that leucine and pyruvate could have independent mechanisms for rescuing cysteine toxicity.

## Discussion

In this study we show that excess of a single amino acid can lead to large-scale changes in cellular amino acid pool and rewire metabolism. As a paradigm, we show that a cysteine-induced change in metabolism is adaptive, and is primarily routed through a translational regulator. We have used genetic, proteomic, transcriptomic and targeted metabolomics approach to get systemic understanding of a cell exposed to metabolic stress (due to excess cysteine).

### Cysteine inhibits translation and rewires amino acid metabolism

Cell maintains homeostasis during excess cysteine by rewiring the amino acid metabolism, and depends on known regulators of amino acid metabolism, Leu3 ^41^ and Stp2 ^42^ for survival. Cysteine increases the levels of lysine and decreases the levels of threonine, and as a consequence the alters the expression of enzymes required for their biosynthesis in opposite directions. Lysine levels have been reported to be high in the yeast strains defective in translational machinery ^43^, thus the levels of lysine may be a marker of translational defect. Cysteine decreases the levels of threonine, and genetic inhibition of threonine biosynthesis further increases the growth sensitivity towards cysteine. The involvement of threonine biosynthesis during high levels of cysteine was also reported in *E.coli* more than three decades ago, where it was suggested that cysteine inhibits homoserine dehydrogenase (Hom6 in yeast) ^44^, acetohydroxyacid synthetase (Ilv6 in yeast) ^45^, threonine deaminase (Ilv1 in yeast) ^46^. Furthermore, cysteine increases the levels of arginine and the enzymes involved in its biosynthesis, and this observation is in agreement with a recent report suggesting that high levels of cysteine increase arginine biosynthesis for polyamine synthesis ^47^.

This study has highlighted the role of amino acid metabolism, and metabolic genotype of strain in governing the cellular growth in high levels of cysteine. We have also shown that cell may need high levels of leucine to survive during stress induced by excess of amino acids. Leucine not only reverses the directionality of the expression of proteins induced by cysteine treatment, but also maintains balanced amino acid levels. Consistent with the reports that leucine assists translation ^48 49^ we also found that leucine rescues cysteine induced translational defect. Thus, this study suggests that the translational defect might be the primary cause of cysteine induced toxicity which further leads to imbalance in amino acid levels.

### NCL1 mediated translation rewires metabolism during high cysteine

It was found that NCL1, a methyltransferase is required for surviving in high levels of cysteine. Appropriate methylation marks have a vital role in the normal development, and loss of NCL1 homolog NSUN2, in mouse and humans, represses global protein synthesis, and leads to neurodevelopmental defects ^50 51 52^. Here, we have shown that in presence of high levels of cysteine, deletion of NCl1 abrogates the correlation between the mRNA and protein levels, and depletes the levels of proteins involved in translational machinery. Most importantly, our results clearly indicate that NCl1 deletion supresses the conversion of Leucine to KIC, and supplementation of KIC and not leucine could rescue the cysteine induced growth defect, highlighting the necessity of conversion of leucine to KIC in this process. Both leucine and KIC, via BAT1 could activate TORC1 independent of EGO1 complex, and it is known that deletion of BAT1 leads to TCA cycle block ^39^. In this study we have shown that expression of BAT1 depends on NCL1, and in ncl1Δ strain, cysteine leads to TCA cycle block, which can be attenuated by KIC and not by leucine.

### Cysteine and pyruvate relation

Our results indicate that cysteine treatment increases the levels of citrate, a known inhibitor of pyruvate kinase ^40^. It has also been reported that cysteine directly inhibits pyruvate kinase and decreases the levels of pyruvate ^53^. The low levels of pyruvate observed in our study could be due to one of these effects. Low pyruvate levels could account for partial growth inhibitory effect of cysteine since addition of pyruvate and leucine together could completely abrogate the growth defect.

Increase in the concentration of cysteine leads to obesity and decreased glucose tolerance ^54^; its prolonged treatment lowers the levels of pyruvate and inhibits glucose induced ATP production in pancreatic cells ^53 55^. Also, in a case control study in the Indian population, we have previously shown that high levels of cysteine is associated with cardiovascular disease ^32^, and interestingly total proteins in CAD patients was shown to be significantly lower as compared to control groups ^56^. Interestingly, high leucine diet has been proposed to be helpful in attenuating heart diseases ^57^. Extrapolating these observations, we speculate that the association of elevated cysteine levels with metabolic disease may be linked through energy metabolism and inhibition of protein translation, which could restore by leucine supplementation.

## Materials and Methods

### Materials

Yeast media constituents including yeast extract, peptone, and dextrose were purchased from Himedia (India), and the amino acids were purchased from Sigma (St. Louis, MO, U.S.A.). IAA (Iodoacetamide), DTT (dithiothreitol), formic acid and ammonium formate was bought from Sigma, and sequencing grade trypsin was procured from Promega. Other MS based reagents and columns were purchased from Sciex as mentioned previously ^58 59^

### Yeast strains, media and growth conditions

The wild-type *S. cerevisiae* strains BY4741 (MATa his3Δ1 leu2Δ0 met15Δ0 ura3Δ0) and S288C used in this study were procured from American Type culture collection (ATCC). The yeast deletion library in the background of BY4741 was obtained from Invitrogen. In S288C background, leu2Δ strain was generated using homologous recombination. Pre-inoculation of all the strains was done in YPD media composed of yeast extract (1%), peptone (2%) and dextrose (2%). The experiments were performed in synthetic complete media (SC media) containing glucose (2%), yeast nitrogen base (0.17%), ammonium chloride (0.5%), adenine (40μg/ml) uracil (20μg/ml) and amino acids ^24^. The amino acids mixture of the SC media were composed of Valine (150μg/ml), threonine (200μg/ml), arginine −HCl (20μg/ml), phenylalanine (50μg/ml), tyrosine (30μg/ml), leucine (60μg/ml), aspartic acid (100μg/ml), lysine (30μg/ml), histidine, (20μg/ml), glutamic acid-monosodium salt (100μg/ml), tryptophan (40μg/ml) ^58 24^. In the growth screen (Fig 1–3), the amino acids were added in the concentration of 10 times higher than mentioned in the SC media. However, for the amino acids which were not present in SC media an arbitrary concentration of 4 mM (close to the average concentration of all the amino acids, 4mM) for Alanine, Glutamine, Isoleucine, Asparagine, Glycine, Serine and Proline, and 1 mM for cysteine was used.

### Growth assay

Yeast cells were pre-inoculated in YPD media overnight in an incubator shaker at 30°C and 200 rpm. The saturated culture was then washed three times with sterile water and re-inoculated at 0.1 O.D in SC media. To study the effect of exogenously added amino acids, cells were treated with excess amino acids and were grown at 30°C with 200 rpm and then growth was measured after 12 hrs. Growth kinetics with cysteine treatment was monitored by withdrawing aliquots at different time points and measuring the turbidity (at 600 nm) using a spectrophotometer (Eppendorf Biophotometer plus, USA).

### Yeast Knock Out

All yeast deletion strains in the background of BY4741, were obtained from Yeast Knock Out Library. The deletion of LEU2 in S288C background was carried with NAT cassette having overhangs for homologous recombination ^60^ using 5’ primer and 3’ primer. The NAT cassete was amplified from pYMN23 plasmid, using Forward primer (AAATGGGGTACCGGTAGTGTTAGACCTGAACAAGGTTTACAGCTTCGTACGCTGCAGGTC) and reverse primer (TTAAGCAAGGATTTTCTTAACTTCTTCGGCGACAGCATCAGCTTCTAATCCGTACTAGAG). The cassette was transformed in S288C strain, using LiAc yeast transformation as described below and selected for NAT resistance.

### Yeast Transformation

Yeast cells were grown to 0.6 OD and then harvested by centrifugation at 2500 g for 5 mins at 30°C. Later, cells were re-suspended in lithium acetate (LiAc) buffer and incubated in shaking for 30 minutes and then centrifuge at 2500 g for 5 mins at 300°C to pellet down the cells. Cells were then re-suspended in 100μl of LiAc buffer, and then template DNA (2 μg) and single stranded carrier DNA (5μg) added. Then, 50% PEG with LiAc buffer was added and vortexed briefly and incubated at 30°C for 90 minutes. Then heat shock was given for 10 min at 42°C. The solution was then centrifuged at 10,000 rpm for 1 min and the supernatant was discarded. The pellet was washed with autoclaved sterile water. The pellet was then resuspended in YPD for 1 hour at 30°C with gentle shaking at 120 rpm and 100-150μl of total sample was plated on YPD-Nat plates.sterile

### Polysome profiling

Yeast cells were grown in presence and absence of 1 mM cysteine for 6 hrs, and then 50μg/ml of cycloheximide was added to the cultured cells and kept in ice for 5 mins to stall the ribosomes. Cells were then processed as described previously ^59^. Briefly, cells were harvested at 2500 g for 5 mins at 4°C, and then lysis was performed with bead beater in lysis buffer (50mM Tris pH −7.5, 150 mM NaCl, 30mM MgCl2, 50μg/ml CHX). RNA content of the samples were normalized by measuring the absorbance at 260nm. Equal amount of RNA (5 OD at 260nm) was then loaded for each of the sample to the 7-47% continuous sucrose gradient (50mM Tris acetate pH-7.5, 250 mM Sodium acetate, 5mM MgCl2, 1mM DTT and 7-47% sucrose) and centrifuged in Beckman SW32Ti rotor at 28000 rpm for 4 hours at 4°C. The samples were then fractionated using ISCO gradient fractionator with detector sensitivity set to 0.5. This fractionator was equipped with a UV absorbance monitor, and thus optical density of the gradients at 254 nm was measured in real time to obtain the profile.

### s^35^ Methionine labelling

De-novo protein translation was measured by the rate of incorporation of S^35^ Methionine in the nascent polypeptides. Yeast cells are grown in the SC media for 0.6-0.8 OD, and then washed with sterile water. Cells were resuspended in SC media without methionine for 10 minutes, and then finally S^35^ methionine and cold methione was added. Then cells were distributed in three groups, one group was kept as control, in other two groups, 1 mM and 2 mM cysteine was added. Then cells were incubated at optimum temperature, and 1 ml aliquots were withdrawn at different time points, 15, 45, 120 and 360 minutes.

Cells were lysed using alkaline lysis method ^61^ and isolated protein was estimated using BCA (Bicinchoninic Acid) method. Equal protein was then loaded on SDS PAGE and stained with Coomassie Blue staining. Incorporation of S^35^ methionine in the isolated proteins was then analyzed by scanning the protein gel using Image safe scanner.

### Protein isolation for MS based proteomics

Cells with and without cysteine treatment were harvested and washed three times with sterile water. As mentioned previously ^58 59^, lysis was done by bead beating method using lysis buffer containing 10mM Tris-HCl (pH 8), 140mM NaCl, 1.5mM MgCl2, 0.5% NP40 and Protease inhibitor cocktail. Isolated protein was then buffer exchanged with 0.5 M Triethyl ammonium bicarbonate (pH 8.5) using a 3 kDa cutoff centrifugal filter (Amicon, Millipore). Protein was then estimated using Bradford’s reagent (Sigma, USA).

### Trypsin digestion and iTRAQ labelling

Buffer exchanged proteins from different groups were digesting using trypsin as mentioned previously ^58 59 62^. Briefly, sixty micrograms of protein from each sample was reduced with 25mM DTT for 30 minutes at 56°C and the cysteine’s were blocked by 55mM IAA at room temperature for 15-20 minutes. This was followed by digestion with modified trypsin added in the ratio of 1:10 ratio (trypsin to protein) and incubated at 37°C for 16 hrs. We used 4-plex and 8-plex labelling strategies. In the 4-plex experiment the proteins digests from cells treated with and without cysteine for 12 hrs were used. However, in the 8-plex experiment protein from Wt and ncl1Δ strains, treated with cysteine and combination of leucine and cysteine for 6 hrs was used. With biological triplicates both the labelling procedures were performed using the manufacturer’s protocol (AB Sciex, Foster City, CA) ^58 59 62^

### 2D nano LC-MS/MS iTRAQ analysis

iTRAQ labelled peptides were first fractionated by cation exchange (SCX) chromatography with SCX Cartridge (5 micron, 300 Å bead from AB Sciex, USA) using a step gradient of increasing concentration of ammonium formate (30 mM, 50 mM, 70 mM, 100 mM, 125 mM, 150 mM, and 250 mM ammonium formate, 30% v/v ACN and 0.1% formic acid; pH= 2.9) ^58 59 62^. The iTRAQ labelled fractions of 4-plex and 8-plex obtained after SCX chromatography were analyzed on quadrupole-TOF hybrid mass (Triple TOF 5600 & 6600, Sciex USA) spectrometer coupled to an Eksigent NanoLC-Ultra 2D plus system.

For each fraction 10μl of sample was loaded onto a trap column (200 μm× 0.5 mm) and desalted at flow rate 2 μL / min for 45 minutes. Then peptides were separated using a nano-C18 column (75 μm × 15 cm) using a gradient method with buffer A (99.9% LC-MS water + 0.1 % formic acid) and buffer B (99.9% acetonitrile + 0.1% formic Acid) as described previously ^58 59 62^. Data was acquired in an Information dependent acquisition ^63^ mode with MS settings as follows: nebulizing gas of 25; a curtain gas of 25; an Ion spray voltage of 2400 V and heater interface temperature of 130°C. TOFMS scan was performed in the mass range of 400-1600 m/z with accumulation time of 250 msec, while the MS/MS product ion scan was performed with mass range of 100-1800 m/z for 70 msec with a total cycle time of approximately 2.05 seconds. Parent ions with more than 150 cps abundance and with a charge state of +2 to + 5 were selected for MS/MS fragmentation. After an MS/MS fragmentation of an ion its mass and isotopes were excluded for fragmentation for 3 seconds. MS/MS spectra were acquired using High sensitivity mode with ‘adjust collision energy when using iTRAQ reagent’ settings ^58 59 62^.

### Database searching and analysis for proteomics experiment

The raw files in the format of “.wiff” containing the spectra of MS and MS/MS were submitted to Protein Pilot v4.0 software (AB Sciex) for protein identification and relative quantification. Paragon Algorithm was used in a “Thorough ID” search mode against the Uniprot-*Saccharomyces cerevisiae* reference dataset (6643 protein sequences). The parameter for identification search includes: IAA as cysteine blocking agent, 4-plex or 8-plex N-terminal iTRAQ labelling, trypsin digestion with two missed cleavages. Global protein level 1% FDR was considered for protein identification ^58 59 62^.

In 4-plex experiment we identified 1041 proteins in all the three biological replicates with a criteria of 1 % global FDR, and more than or equal to two unique peptides (Supp. Table 6). Proteins were considered to be differentially expressed, if the fold change with respect to control was ≥ 2 or ≤ 0.5 in all the three replicates. With the similar criteria of identification, 543 proteins were identified in all the three biological replicates of 8-plex experiment (Supp. Table 7). Based on fact that fold change of protein expression is generally underestimated in 8-plex iTRAQ experiment ^64 63^, and the narrow range of fold change in the present data set, a fold change cutoff of ±1.15 was considered ^65 66^. Thus, proteins were considered to be differentially expressed if the average fold change was ≤0.85 or ≥1.15, with a similar trend in atleast two replicates.

### Amino acid profiling

Intercellular amino acids were extracted using boiling water lysis method ^67 68^. Briefly, cells (5 OD) were washed three times with MilliQ water, and then reconstituted in 500μl sterile water, and boiled for 10 minutes. Cell debris was removed by centrifugation for 5 minutes at 6000 rpm. Amino acids were then measured by automated pre-column o-Phthaldialdehyde derivatization using HPLC (1290 Infinity LC system, Agilent Technologies Pvt. Ltd). The mobile phases consisted of 10 mM Na2HPO4 and 10mM Na2B4O7 (Buffer A) and Acetonitrile-Methanol-H2O in the ratio of 45:45:10 by volume (Buffer B). In the automated pre-conditional method, 1μL from sample was mixed with 2.5μL of borate buffer (Agilent P/N 5061-3339), followed by addition of 0.5μL of OPA dye (Agilent P/N 5061-3335). Finally, 32μL of Diluent (1 ml Buffer A + 15μL phosphoric acid) was added and 6 μL of mixture was loaded, and separated using Poroshell HPH-C18 column (2.7μm, 3.0×100mm). Amino acids were eluted from the column at a rate of 0.9 ml/min for 14.5 min, using a linear gradient of 2% to 57% of Buffer B for 13.5 min. The eluted amino acids were detected by the fluorescent detector. (Ex-340 nm, Em-450 nm).

### Genetic screen for growth sensitivity

A commercially available S. cerevisiae library of approx. 4500 non-essential genes was used. All the deletion strains in this library contain a specific gene deleted, with KanMX resistance cassette. Primary culture was obtained by growing these strains overnight in 200 μl of YPD media containing G418, in 96-well format. Then, 5 μl of this culture was transformed to another 96-deep well plate, containing 400 μl of SC media, with and without 1 mM cysteine. The strains were then grown at 30°C in a shaking incubator (Thermo scientific, USA), with 200 rpm, for 12 hours, and then, cell density was measured using multimode reader.

### RNA isolation

RNA was isolated from both Wt and ncl1Δ, treated with and without 1 mM cysteine for 6 hours. Acid-phenol based method was used to isolate total RNA and then purified using RNAeasy mini kit (Qiagen,USA) after a DNAse treatment (TURBO™ DNase, Ambion) following the manufacturer instruction ^69^. The RNA was examined on ethidium bromide-stained agarose gel to confirm its integrity.

### Library generation and mapping of sequencing reads

Library was prepared with 700ng RNA of each sample using Truseq RNA sample prep kit v2 (stranded mRNA LT kit) according to the manufacturer’s instructions (Illumina Inc., USA) ^70^. AMPure XP beads (Agencourt) were used to purify Adaptor-ligated fragments, and then amplified (12-14 cyclles), purified and finally measured using Qubit instrument (Invitrogen). Then, using BioAnalyzer DNA1000 LabChip (Agilent Technologies) the average fragment size of the libraries was determined. And finally the diluted libraries were multiplexed and loaded on HiSeq Flow Cell v3 (Illumina Inc., USA). Sequencing runs were performed on a HiSeq 2000 Illumina platform using TruSeq SBS Kit v3 (Illumina Inc., USA) for cluster generation as per manufacturer’s protocol.

Sequencing reads with phred quality score equal or greater than 30 were taken for analysis. Read sequences were trimmed using Trimmomatic (v0.43), and then aligned to the transcriptome of *S. cerevisiae* strain S288c as available from ENSEMBL using Kallisto (v0.36) software. Gene expression levels were then estimated as TPM (Transcripts per million) values.

### Targeted metabolomics

Metabolites for MS based targeted metabolomics were extracted using cold methanol method. Cells (50 OD) from different treatment groups were washed three times with Sterile water, and then quenched with chilled ethanol (kept at −80 °C), and followed by bead beating using acid washed glass beads. The suspension was then transferred to a fresh tube and centrifuge at 13000 rpm at 4 °C for 10 minutes. Supernatant was then vacuum dried, and then reconstituted in 50μl of 50% methanol. The reconstituted mixture was centrifuged at 10000 rpm for 10 minutes, and 5 μl was injected for LC-MS/MS analysis.

The data were acquired using a Sciex Exion LCTM analytical UHPLC system coupled with a triple quadrupole hybrid ion trap mass spectrometer (QTrap 6500, Sciex) in a negative mode. Samples were loaded onto an Acquity UPLC BEH HILIC (1.7μm, 2.1×100 mm) column., with a flow rate of 0.3 mL/min. The mobile phases consisted of 10mM ammonium acetate and 0.1% formic acid (Buffer A) and 95% Acetonitrile with 5mM ammonium acetate and 0.2% formic acid (Buffer B). The linear mobile phase was applied from 95% to 20 % of Buffer A. The gradient program was employed as follow: 95% buffer B for 1.5 minutes, 80% to 50% buffer B in next 0.5 minute, followed by 50% buffer B for next 2 minutes, and then decreased to 20% buffer B in next 50 seconds, 20% buffer B for next 2 minutes and finally again 95% buffer B for next 4 minutes. Data was acquired using three biological triplicates with three technical replicate for each run. A list of measured metabolites and their optimized parameters are given in the table (Supp. Table 8). Relative quantification was performed using MultiQuantTM software v.2.1 (Sciex)

Pyruvate measurement was done using syringe pump based direct infusion method. Lyophilised samples were reconstituted in 250ul of 50% methanol. Harvard syringe pump coupled with 6500 QTRAP in negative mode was used. Infusion parameters were set to as follow: ion spray voltage: −4500 V, CUR gas: 35, GS1: 20, GS2: 10, CAD: high, DP: −60V, CE: −38V, flow rate: 7ul/min. MRM transition used for pyruvate was 87/43 and peak intensities obtained was used for relative quantification.

### Data availability

Transcriptomic data generated in the manuscript has been submitted to SRA database under the accession number PRJNA514239. The processed data files of proteomics data has been provided in the supplementary section of this manuscript, and the raw files are available from corresponding author on request.

## Acknowledgement

We acknowledge the financial support from the Council of Scientific and Industrial Research (CSIR), India. The study was funded under the project titled ‘CARDIOMED: Centre for Cardiovascular and Metabolic Disease Research (BSC0122)’. R. C. acknowledges the Junior Research Fellowship from UGC. We thank Dr. Shuvadeep Maity and Dr. Tryambak Basak, for their contribution in standardizing yeast and proteomics experiments, respectively. We are grateful to Asmita Ghosh and Sarada Das for transcriptomic experiments. We thank Dr. Vignesh Kumar for his help in standardizing polysome profiling experiment. We are also thankful to Zeeshan Hamid and all master students for being valuable part of this project.

## Author Contribution

AB, KC and SSG designed the project. AB, RC and GA were involved in yeast experiments. AB and RC were involved in proteomics experiment and LC based amino acid measurement. KA and RC standardized and performed LC-MS based targeted metabolomics experiments. AB, KC and SSG were involved in data analysis and representation of the results. AB, KC and SSG wrote the manuscript with inputs from all authors. KC and SSG supervised the work. The final version of manuscript was read and approved by all the authors.

## Supplementary Figure legends

Figure S1: **Dose dependent growth inhibition induced by cysteine**.

A. Growth kinetics of BY4741 strain during different concentration of cysteine.

Figure S2: **Cysteine induced effect on protein translation**

A. Functional classification based on biological function of differentially expressed proteins. Protein was isolated from cells treated with cysteine for 6 hrs and 12 hrs, and iTRAQ based proteomics was performed. Differentially expressed proteins at each time interval were classified based on biological function using DAVID, and top ten enriched pathways with p-value ≤ 0.05 are plotted. The above panels are for proteins upregulated by cysteine at 12 hrs (left panel) and 6 hr (right panel) respectively, and similarly below panels are for down-regulated proteins.

B. S^35^-Methionine incorporation assay in Δmet6 deleted strain (cannot metabolize homocysteine to methionine). The upper panel represents the coomassie stain of total protein, and the lower panel represents the autoradiogram of the labeled proteins.

Figure S3. **NCL1 mediated regulation of proteome during cysteine treatment**

A. The bar graph represents the biological pathways enriched from proteins differentially expressed in ncl1Δ strain during cysteine treatment, with respect to their expression in cysteine treated Wt cells. Left and right panel represents the biological pathways up-regulated and down-regulated, respectively in ncl1Δ.

B. Box plot represents the relative expression of proteins during co-treatment of leucine and cysteine in Wt and ncl1Δ, for proteins which were downregulated by cysteine in Wt.

### Supplementary Table

Table 1: Differentially expressed proteins due to treatment of cysteine for 12 hrs

Table 2: Differentially expressed proteins due to cysteine treatment for 6hrs, and their status during co-treatment of leucine and cysteine.

Table 3: This table includes the percentage growth of cysteine sensitive deletion strains in comparison to wild type strain during cysteine treatment.

Table 4: List of proteins differentially expressed in ncl1Δ strain during cysteine treatment with respect to wild type strain

Table 5. Relative expression of genes at RNA and proteins levels during cysteine treatment in Wt and ncl1Δ strain.

Table 6: Proteomic prolife of all proteins identified in 4-plex experiment from wild type cells treated with cysteine for 12 hrs.

Table 7. Proteomic prolife of all proteins identified in 8-plex experiment from ncl1Δ and Wt strain treated with cysteine and combination of leucine and cysteine for 6 hrs.

Table 8: Optimized parameters for targeted metabolomics

