## Supplementary figures and images for "Ncl1 mediated metabolic rewiring critical during metabolic stress"

### Supplementary Figure 1

Supp. Figure 1

A. Growth kinetics in presence of different concentration of cysteine

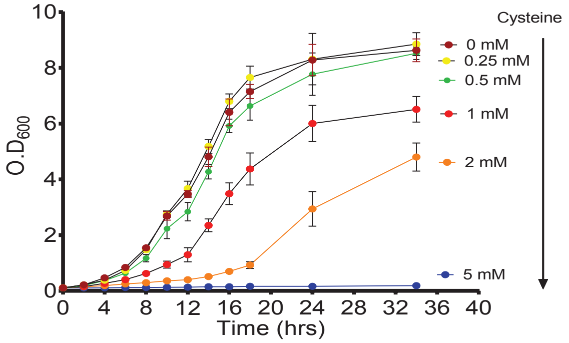
