## Supplementary Figure 2 for "Ncl1 mediated metabolic rewiring critical during metabolic stress"

Supp. Figure 2

A. Biological process enrichement of differentially expressed proteins in 6 and 12 hrs of cysteine treatment

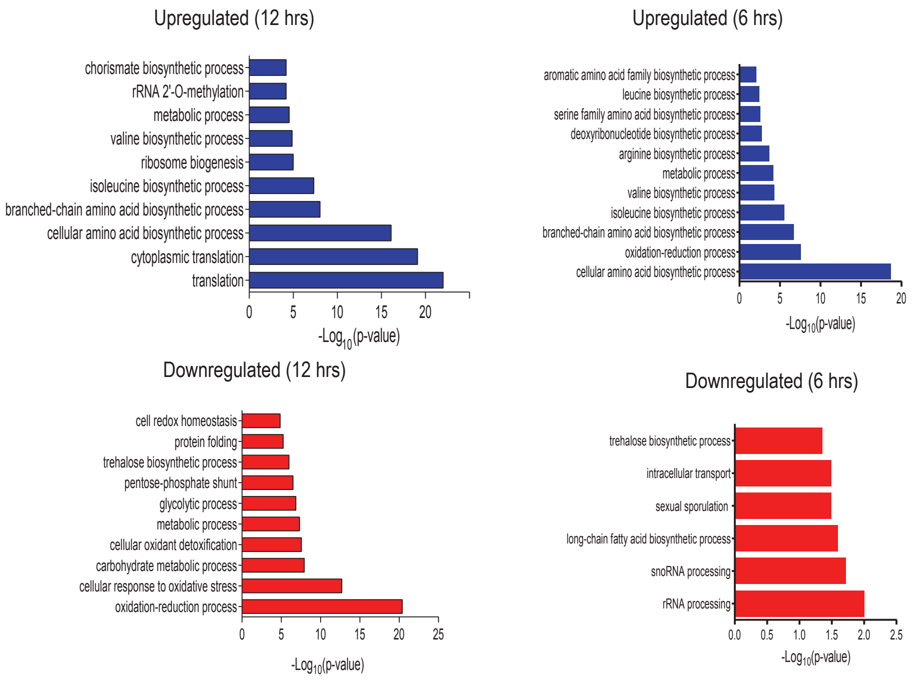


B. Radiolabelled methionine incorporation assay for met6Δ strain during cysteine treatment

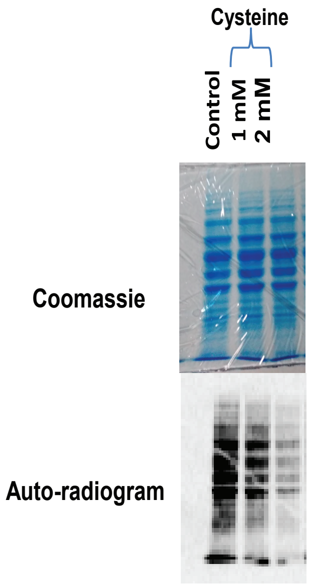
