## Supplemental Figure 3 for "Ncl1 mediated metabolic rewiring critical during metabolic stress"

Supp. Figure 3

A. Biological enrichment of proteins upregulated (right panel) and downregulated by cysteine in ncl1Δ cells with respect to Wt.

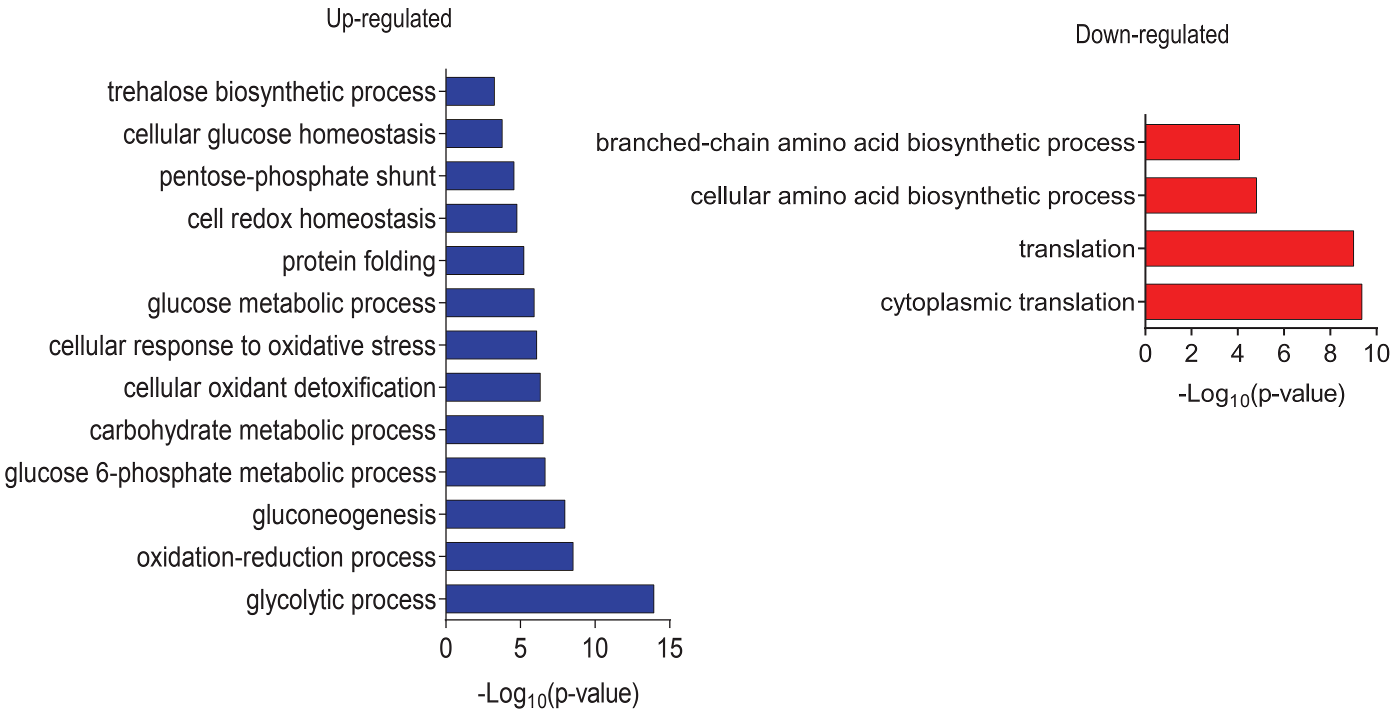


B. Leucine reverts the expression of proteins downregulated by cysteine in Wt, but does not alter their expression in ncl1Δ

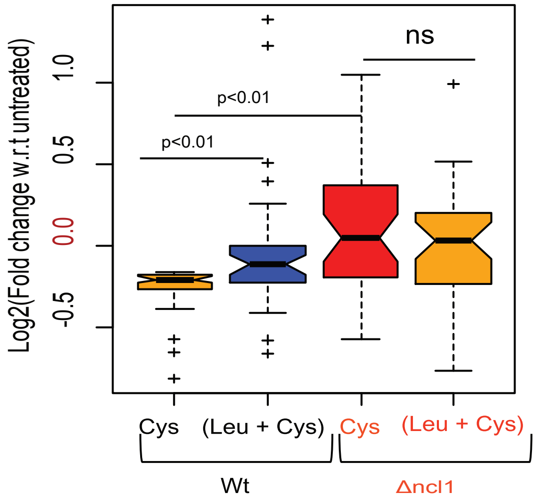
